## Supplementary material for "SARS-CoV-2 surveillance between 2020 and 2021 of all mammalian species in two Flemish zoos (Antwerp Zoo and Planckendael Zoo)"

**Table S1. *Details of serum samples screened.***

|  | **N tested** | |
| --- | --- | --- |
| **Order and species tested** | **N collected before 2020** | **N collected from 2020 onwards** |
| **Artiodactyla** |  |  |
| *Addax nasomaculatus* |  | 5 |
| *Camelus bactrianus* |  | 1 |
| *Equus grevyi* | 1 | 1 |
| *Equus zebra hartmannae* | 2 | 2 |
| *Gazella leptoceros* | 1 | 1 |
| *Giraffa camelopardalis antiquorum* | 2 | 2 |
| *Nanger dama mhorr* |  | 1 |
| *Okapia johnstoni* | 1 | 1 |
| *Oryx dammah* |  | 1 |
| *Tragelaphus eurycerus isaaci* |  | 1 |
| *Vicugna pacos* |  | 1 |
| *Vicugna vicugna* |  | 1 |
| **Carnivora** |  |  |
| *Panthera leo*  *Panthera leo persica* | 1  1 | 1  1 |
| *Speothos venaticus* |  | 1 |
| **Diprotodontia** |  |  |
| *Dendrolagus goodfellowi buergersi* |  | 1 |
| *Macropus giganteus giganteus* | 1 | 1 |
| *Macropus rufus* |  | 1 |
| **Perissodactyla** |  |  |
| *Tapirus indicus* |  | 1 |
| **Macroscelidea** |  |  |
| *Rhynchocyon petersi* |  | 1 |
| **Primates** |  |  |
| *Pan paniscus* | 1 | 3 |
| *Pan troglodytes* | 1 | 3 |
| *Varecia rubra* | 1 | 1 |
| **Proboscidea** |  |  |
| *Elephas maximus* | 1 | 1 |
| **Rodentia** |  |  |
| *Hystrix africaeaustralis* |  | 1 |
| *Phloeomys pallidus* |  | 1 |
| **TOTAL** | 14 | 36 |

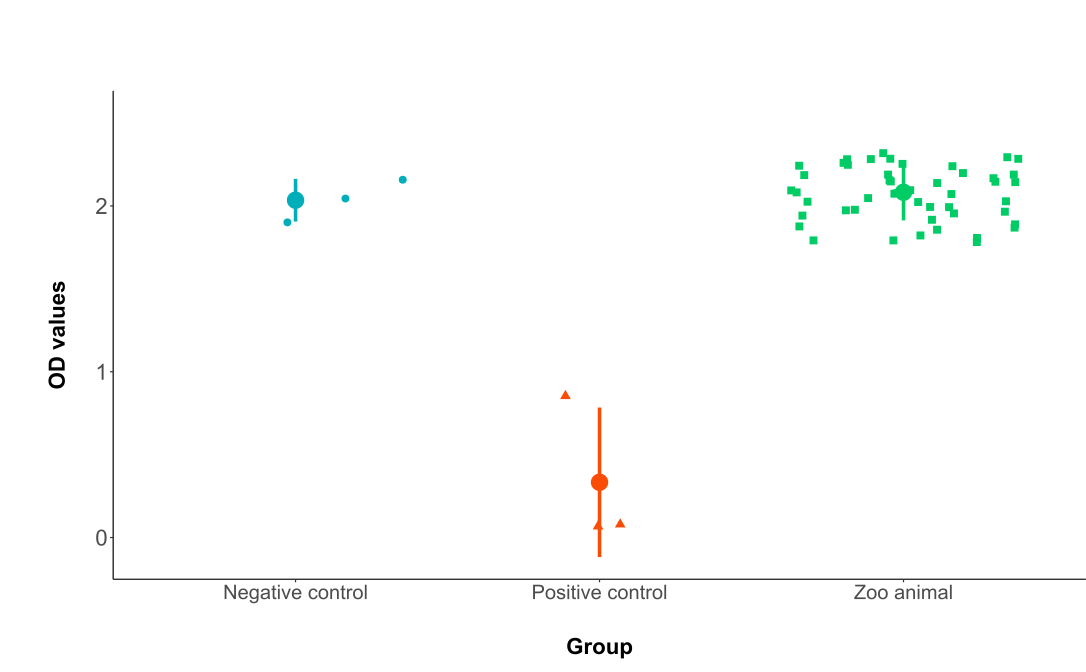

***Figure S1*.** Serological screening of serum samples from various mammal zoo species. OD values of the negative controls, positive controls and zoo animals’ samples are shown respectively in blue, red and green. Mean and SD (Standard Deviation) values are represented for each group.

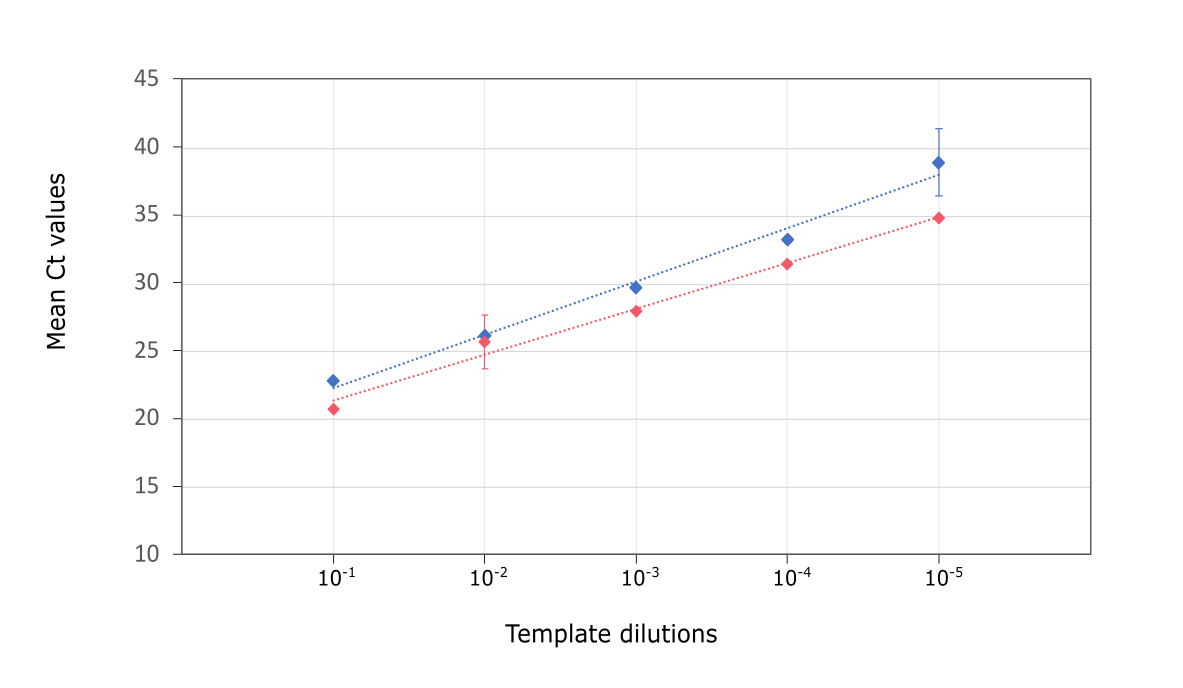

***Figure S2***. Graph of mean PCR cycle threshold (Ct) vs. ten-fold dilutions of SARS-CoV-2 template for each PCR system target. Blue: Pan-CoV PCR system; Red: CDC specific SARS-CoV-2 PCR. Each square represents a serial dilution from the initial template with respective standard deviation based on triplicate assay. The dashed lines represent the trendline for each values set. The linear equation of the trendline for the Pan-CoV PCR system is 3.92 + 18.32 with R²= 0.99. The linear equation for the CDC specific SARS-CoV-2 PCR is 3.38 + 17.92 with R²= 0.99.
